# Identification of *Cryptococcus neoformans* capsule-associated protein antigens that mediate B cell-dependent vaccine protection in mice and elicit antibody responses in humans with cryptococcosis

**DOI:** 10.64898/2026.10.28.762512

**Authors:** Maureen M. Hester, Diana Carlson, Lorena V. N. Oliveira, Zachary Hastings, Satya Sahu, Jessica Hargarten, Seher Anjum, Gary R. Ostroff, Peter R. Williamson, Charles A. Specht, Stuart M. Levitz

## Abstract

Infection with the encapsulated fungal pathogen *Cryptococcus neoformans* causes substantial global disease burden in persons with compromised T cell function. Previously, we demonstrated that vaccines containing adjuvanted recombinant cryptococcal proteins protected mice against lethal infection by mechanisms dependent on CD4^+^ T cells. For the present study, we hypothesized that vaccines containing protein antigens naturally located on the surface of the cryptococcal capsule would contribute protective B cell responses. We found that B cell-deficient mice exhibited reduced survival following vaccination with any of four capsule-associated antigens: Blp4, Cpd1, Lhc1, or Krp1. To further elucidate the role of antibodies in protection, we tested serum from mice vaccinated with a live-attenuated strain (*cda123*) and then challenged with a virulent *C. neoformans* strain for IgG responses to a panel of 26 cryptococcal recombinant proteins. Strong reactivity was detected against Blp4, which we focused on because it is capsule-associated and protects mice when administered in adjuvanted subunit vaccines, by mechanisms partly dependent on B cells. Blp4 antibodies were consistently present in surviving mice, including those vaccinated with unrelated proteins or those challenged with diverse clinical isolates. Additionally, serum from Blp4-vaccinated mice promoted macrophage opsonophagocytosis of *C. neoformans*. Finally, IgG antibodies against Blp4 were detected in 20 of 21 serum samples from patients who survived cryptococcosis, compared with only 4 of 14 samples from healthy controls. Together, these data identify Blp4 as an antigen that elicits protective B cell responses and is a strong candidate for inclusion in subunit cryptococcal vaccines.

## Introduction

Fungal infections cause an estimated 2.5 million deaths each year, and this burden is heavily skewed to resource-limited regions of the world [1]. Despite this toll, there are currently no licensed fungal vaccines. One such devastating disease is cryptococcal meningitis (CM), occurring predominantly in populations with compromised CD4^+^ T cell function following infection with *Cryptococcus neoformans* and its sister species *C. gattii*. The cause of nearly 120,000 deaths annually, CM is responsible for approximately 19% of AIDS-related mortalities [2]. Although globally distributed, rates of CM are highest in sub-Saharan Africa where upwards of 25 million inhabitants are fighting AIDS [2]. The World Health Organization categorizes *C. neoformans* and *C. gattii* as critical- and medium-priority pathogens, respectively, for which treatability and management problems currently exist [3]. Successful development of a cryptococcal vaccine is paramount to addressing the increasingly urgent need to protect vulnerable populations, including immunosuppressed individuals and those living in areas where pathogenic *Cryptococcus* species are endemic.

While cryptococcal vaccines have not advanced to clinical trials, vaccines show promise in preclinical studies of experimental cryptococcosis in mice (reviewed in [4]). Our lab has demonstrated success in protecting mice against an otherwise lethal *C. neoformans* infection using both live-attenuated and subunit vaccines. The live-attenuated vaccine we utilize is an avirulent, chitosan-deficient *C. neoformans* mutant, *cda1*Δ*cda2*Δ*cda3*Δ (*cda123*), generated through the deletion of three chitin deacetylase (*CDA*) genes [5]. *cda123* is rapidly cleared from the lungs of mice and selectively enriches for protective T helper type 1 (Th1) cell populations [6], providing robust protection against reference and clinical strains of *C. neoformans* [6-8]. We have also individually tested 23 subunit vaccines, each containing a recombinant cryptococcal protein delivered in glucan particles (GPs) or adjuvanted with the liposomal Mincle agonist Cationic Adjuvant Formulation 01 (CAF01); 12 of these vaccines significantly protected C57BL/6 or BALB/c mice [8-14]. Using genetically deficient mice and mice depleted of specific cell types with monoclonal antibodies, we have extensively studied the immunological mechanisms underpinning protection following vaccination with live-attenuated *cda123*, as well as with subunit vaccines containing the protein antigens Cda1 and Cda2. We found that while B cells and CD8^+^ T cells are dispensable, CD4^+^ T cells and interferon-gamma (IFNγ) are required for protection when using either *cda123* or subunit vaccines [7, 12].

Protection against *C. neoformans* is classically dependent on CD4⁺ T cell-mediated Th1-type immunity (e.g., IFNγ, tumor necrosis factor α), as supported by both human disease and murine models in which CD4⁺ T cells are essential for fungal control and Th1 responses are protective [4, 15]. This creates a central paradox: Individuals most in need of a vaccine are those with impaired CD4⁺ T cell function. However, the immune mechanisms required for protection against natural infection may differ from those sufficient for vaccine-mediated immunity. Emerging evidence indicates that protective responses can engage alternative pathways, such as by antibody-mediated mechanisms, even in CD4⁺-deficient settings. Indeed, antibody-mediated immunity contributes to host defenses against cryptococcosis, as evidenced by associations between FcγR3A polymorphisms and susceptibility to AIDS-associated cryptococcal meningitis [16], and by the presence of anti-cryptococcal antibodies in humans [17, 18]. As such, an ideal vaccine candidate would induce a protective antibody response in addition to stimulating CD4^+^ T cell responses.

The carbohydrate capsule of *Cryptococcus* is the organism’s main virulence factor. Vaccines designed to elicit protective antibody responses to glucuronoxylomannan (GXM), the major component of capsule, have met with mixed success in preclinical models (reviewed in [19]). Protection may be impaired because antibodies form immune complexes with GXM shed by *Cryptococcus* before reaching fungal cells [20]. Recently, three cryptococcal proteins were shown to co-purify with the outer layer of the cryptococcal capsule [21]. These antigens, Barwin-like protein 4 (Blp4), Carboxypeptidase 1 (Cpd1), and Lactonohydrolase 1 (Lhc1), protect mice when formulated as GP-vaccines [10, 11]. In the present study, we examined the antibody response to capsule-associated protein antigens during mouse and human infection, including their contribution to vaccine-mediated protection.

## Results

We previously demonstrated the efficacy of recombinant cryptococcal antigens delivered in glucan particles in protecting mice against an otherwise lethal *C. neoformans* KN99 challenge [9-11]. To determine the role of antibodies in vaccine-mediated protection, we compared the survival of C57BL/6 and muMT mice. muMT mice are on the C57BL/6 background but lack mature B cells and have antibody levels over 100-fold lower than their wild-type counterparts [22]. We found GP-vaccines containing Cda1 or Cda2 protected C57BL/6 and muMT mice from cryptococcal challenge to a similar extent [12]. However, while Cda1 and Cda2 are found in the cell wall and in extracellular vesicles, they are not exposed on the capsular surface [23, 24]. It has been shown that other cryptococcal vaccine protein antigens, including Blp4, Cpd1, Lhc1, and Kelch repeat-containing protein 1 (Krp1), co-purify with the outer layers of capsule [21] or affect capsular architecture [25]. Therefore, we hypothesized that antibodies could contribute to the protection mediated by these capsule-associated cryptococcal antigens.

To test this hypothesis, we vaccinated (subcutaneous prime followed by one boost three weeks later) muMT and C57BL/6 mice with Cda1, Cda2, Blp4, Cpd1Δ, Lhc1, or Krp1, each adjuvanted with CAF01. Note that Cpd1Δ refers to the Cpd1 recombinant protein which was deleted of the portion of the sequence with human homology [14]. Two weeks after the booster, mice received an orotracheal challenge with *C. neoformans* KN99 and were followed daily for survival **(Figure 1A**). Unvaccinated control mice all died within 30 days (**Figure 1B**). Consistent with what was shown previously with GP-vaccinations [12], there was no significant difference in survival between muMT and C57BL/6 mice following vaccination with either Cda1 or Cda2 adjuvanted with CAF01 (**Figure 1C, 1D**). However, vaccines containing the capsule-associated antigens Blp4, Cpd1Δ, Lhc1, and Krp1 did not protect muMT mice as well as they did C57BL/6 mice (**Figure 1E-H**). The survival differences were statistically significant for Blp4 and Krp1, but only trends towards significance (p<0.1) for Cpd1Δ and Lhc1.

**Figure 1:**
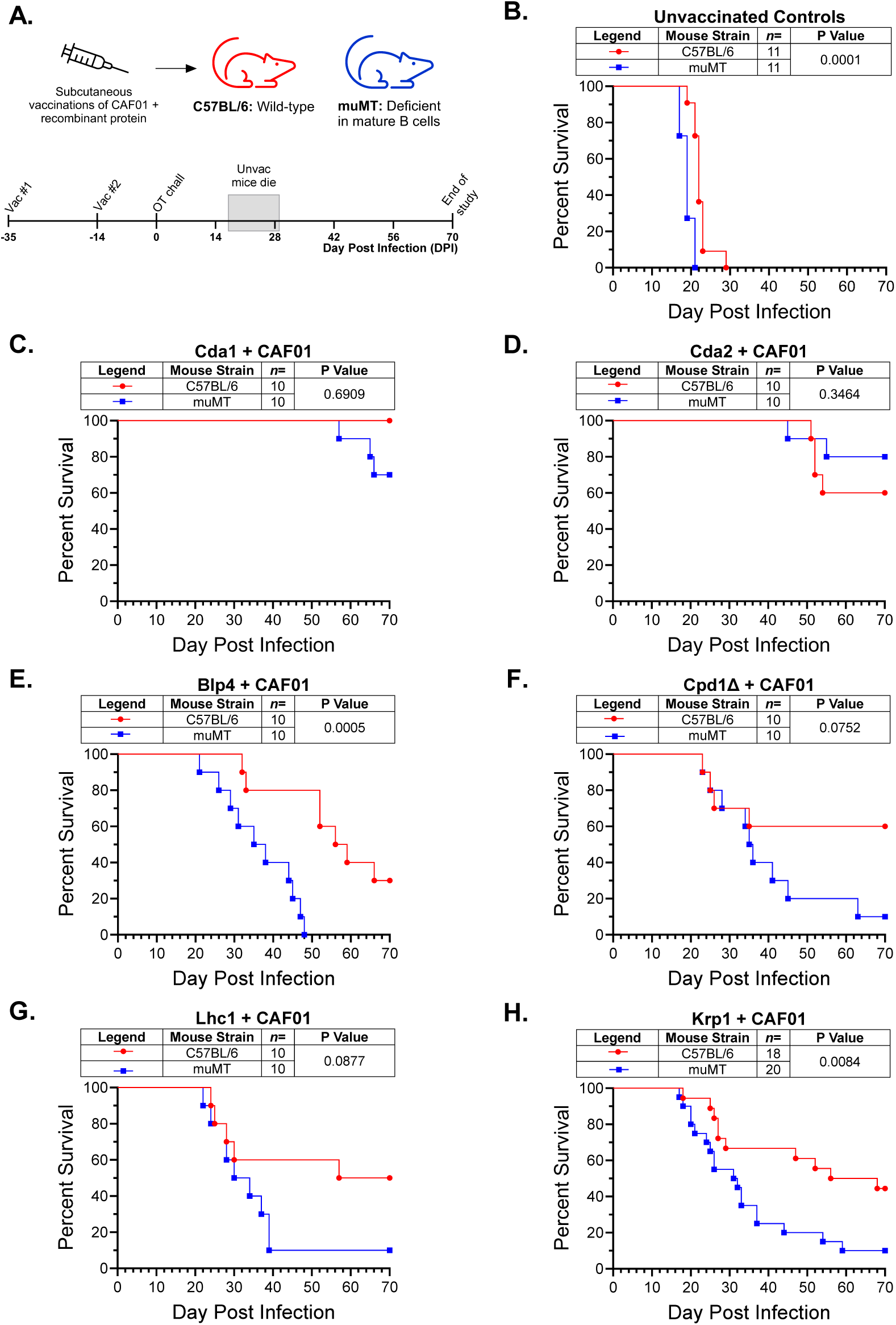
Contribution of B cells to protection by CAF01-adjuvanted vaccines. (A) Schematic of the experimental design. Wild-type C57BL/6 mice and B cell-deficient (muMT) mice on the C57BL/6 background received a prime followed by a boost vaccination three weeks later. Each vaccination consisted of two injections of 5 μg of protein. Two weeks after the boost, mice received an orotracheal challenge with 1 x 10^4^ CFU of KN99 and followed for 70 days for survival. Unvaccinated mice died between 16 and 29 DPI. (B) Survival curves of unvaccinated controls. (C-H) Survival curves following the indicated subcutaneous CAF01-adjuvanted vaccine. Kaplan-Meier survival curves were compared using the Mantel-Cox, log-rank test.

Our results with CAF01-adjuvanted Cda1 and Cda2 parallel what we previously observed using GP-vaccines containing Cda1 or Cda2 in that survival curves were similar when comparing muMT and wild-type C57BL/6 mice (**Figure 1C and D** and [12]). We next examined whether there was a survival difference in the mouse strains when GP-based vaccines containing Blp4 and Cpd1 were used (**Supplemental Figure 1**). We found muMT mice vaccinated with either Blp4 or Cpd1 died significantly sooner than similarly vaccinated C57BL/6 mice following orotracheal challenge with *C. neoformans* (**Supplementary Figure 1**).

Taken together, the data presented thus far demonstrate a protective role for B cells following vaccination with formulations containing capsule-associated protein antigens. We next sought to determine the contribution of antibody to the observed phenotype. Previously, we demonstrated that immunization with GP-Cda1 and GP-Cda2 elicited serum IgG to Cda1 and Cda2, respectively; additionally, we showed that muMT mice vaccinated with GP-Cda2 were unable to generate serum IgG to Cda2, unlike vaccinated C57BL/6 mice [11]. C57BL/6 mice immunized with single antigen CAF01-adjuvanted formulations containing Blp4, Cpd1, Krp1, Lhc1, Cda1, or Cda2 developed IgG antibody responses to the immunization antigen, as measured by Western blotting (**Supplementary Figure 2**). Thus, immunization generates an antibody response to all the cryptococcal proteins tested, regardless of whether B cells were observed to contribute to protection.

Having identified a potential protective role for B cells when vaccinating with formulations containing capsular antigens, we questioned whether we could identify additional B cell-associated antigens by testing mouse sera for dominant antibody responses to other cryptococcal proteins. Through our previous work designing and testing cryptococcal subunit vaccines *in vivo*, we have amassed a library of 26 recombinant cryptococcal proteins purified from *E. coli* (**Table 2**). By dot blotting, we screened pooled serum collected from C57BL/6 and BALB/c mice for reactivity of each strain to these 26 proteins (**Figure 2**). KN99 whole cell protein extract served as a positive control while negative controls included buffer only and recombinant ovalbumin (Ova) produced in *E. coli*. Three conditions were studied: 1. unvaccinated and unchallenged (naïve), 2. vaccinated with *cda123* but unchallenged, and 3. *cda123*-vaccinated mice surviving *C. neoformans* challenge with KN99. Reactivity was not seen in serum from naïve mice (**Figure 2, top**). Mice vaccinated with live *cda123* produced serum IgG reactive with saccharopepsin (Sacch) in both strains of mice (**Figure 2, middle**). Saccharopepsin is an aspartic protease that, when adjuvanted in GPs, protects BALB/c mice (but not C57BL/6 mice) against KN99 challenge [10]. Additionally, the C57BL/6 mouse serum had reactivity to Cerevisin (Cerev) and CNAG_02225. For the vaccinated and challenged mice, the notable finding was the appearance of serum IgG antibody reactive with Blp4 (**Figure 2, bottom**).

**Figure 2:**
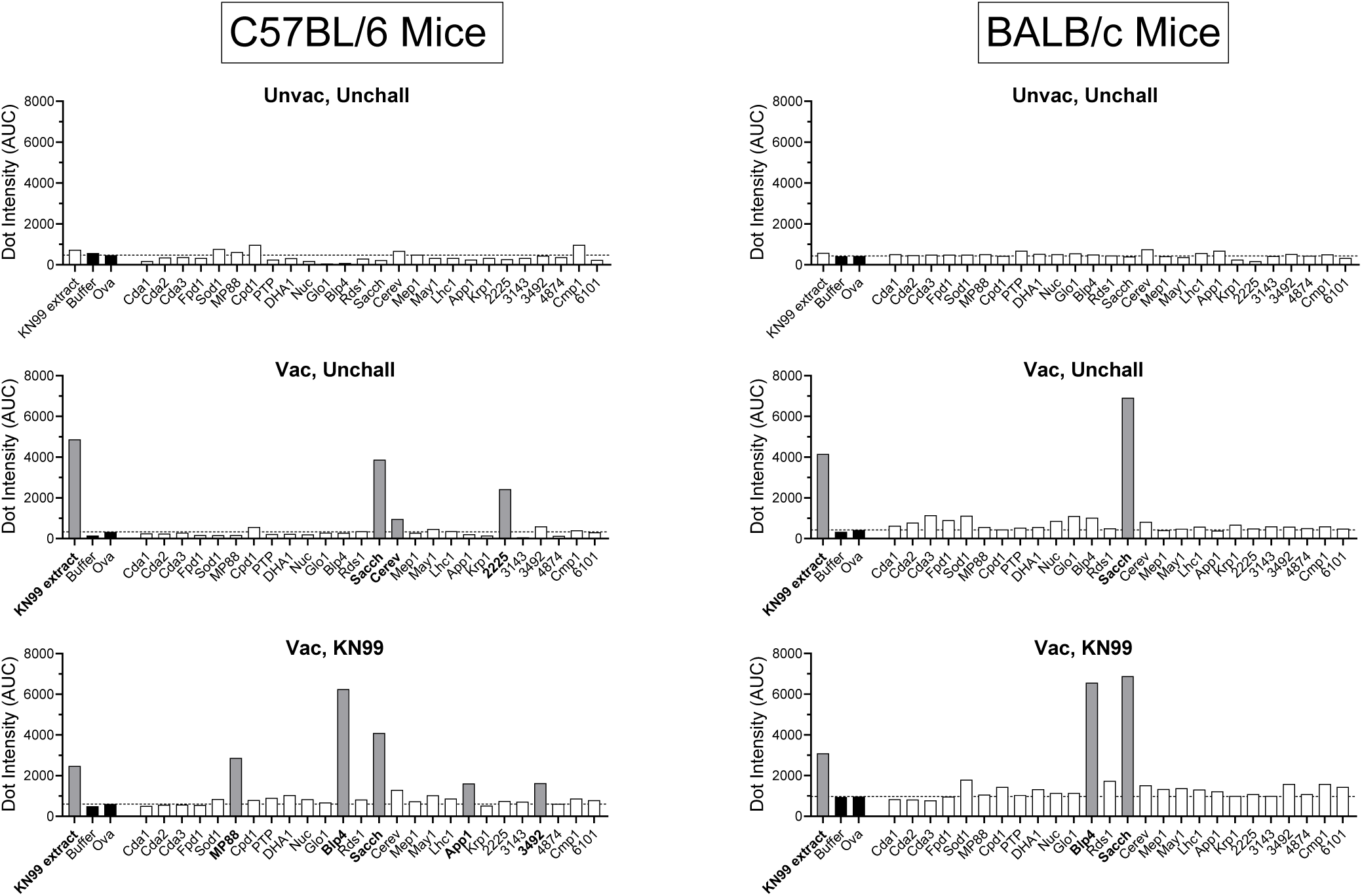
Antibody to cryptococcal antigens in *cda123*-vaccinated mice. Individual recombinant proteins (2 μg), and KN99 extract (20 μg, grown in YPD), were spotted on nitrocellulose. Pooled serum from each treatment group (n=2-6 mice) was used to probe each blot for C57BL/6 mice (left) or BALB/c mice (right). The top panels (Unvac, Unchall) show dots blots using sera from naïve mice. The middle panels (Vac, Unchall) show dots blots using sera from mice that were given an orotracheal vaccination with live *cda123*, followed by two biweekly subcutaneous boosts. Serum was collected two weeks following the final boost. The bottom panels (Vac, KN99) show dot blots using sera from mice that were vaccinated as above and then challenged with *C. neoformans* strain *KN99*. Serum was collected from mice at 70 DPI. Diluted mouse serum was used as the primary antibody, and alkaline phosphatase-conjugated IgG was used as the secondary antibody. Dot intensity was measured using ImageJ, and the resulting area under the curve (AUC) was graphed. Negative controls (buffer only, Buffer; and recombinant ovalbumin, Ova) are denoted by black bars. The dotted horizontal lines indicate the dot intensity of the Ova control. Dot intensities 2.5X higher than that of Ova were considered positive and indicated by grey bars.

The dot blots in **Figure 2** were obtained using pooled mouse sera. When sera from individual vaccinated and challenged mice were studied, IgG antibody to Sacch and Blp4 was seen in each sample (**Supplementary Figure 3A**). The specificity of the dot blots was confirmed by Western blotting which demonstrated reactivity to protein of the expected molecular size (**Supplementary Figure 3B**). The above studies were with mice vaccinated with *cda123*. Similar findings were obtained with mice vaccinated with the cryptococcal antigens Lhc1 and Cda2 in GPs. These mice lacked serum antibody to Blp4 and Sacch before cryptococcal challenge, but the vaccinated mice that survived challenge with *C. neoformans* developed serum IgG reactive to both of those proteins (**Supplementary Figure 4**).

Based on the above results, we chose to focus on Blp4 for additional studies for four main reasons: 1. Blp4 is associated with the cryptococcal capsule, 2. Vaccination with Blp4 confers protection against KN99 challenge in both BALB/c and C57BL/6 mice, 3. Protection conferred by Blp4-containing vaccines is diminished in mice lacking mature B cells, and 4. IgG to Blp4 develops naturally following cryptococcal challenge. Although antibody to Sacch was also identified in the dot blot screen as a dominant response in surviving mice, it is not a protective vaccine antigen in C57BL/6 mice [10], and it is not known to be associated with the fungal capsule. Additionally, while vaccines containing the capsule-associated antigens Cpd1, Lhc1, and Krp1 did not protect muMT mice as well as wild-type, *cda123*-vaccinated and infected mice did not develop a measurable IgG antibody response to any of these antigens.

Blp4 contains a conserved Barwin-like domain which is characteristic of the Blp family of proteins in *Cryptococcus*, consisting of Blp1 through Blp6 [26]. Each of the proteins in this family contain a signal peptide and conserved Barwin-like domain but are otherwise unique in sequence. We previously demonstrated that immunologic cross-reactivity between members of the Cda family contributes to the protection afforded by vaccination with Cda proteins [11]. We therefore sought to determine whether the antibody response was specific to Blp4 or cross-reactivity with other Blp family proteins. To gauge specificity, we utilized a cryptococcal strain with a deletion in *BLP4* (*blp4*Δ). Importantly, *blp4*Δ is a virulent strain, and all unvaccinated mice died by 26 days post infection (**Figure 3A**). Additionally, there was no significant survival difference between groups receiving KN99 or *blp4*Δ challenges, regardless of having been vaccinated with either *cda123* or with a mix of Cda1 and Cda2 protein (Cda1+2). The lung weight and fungal burden in the lungs, brains, and spleens were also similar in mice challenged with KN99 and *blp4*Δ for both sets of vaccinated mice. (**Supplementary Figure 5**).

**Figure 3:**
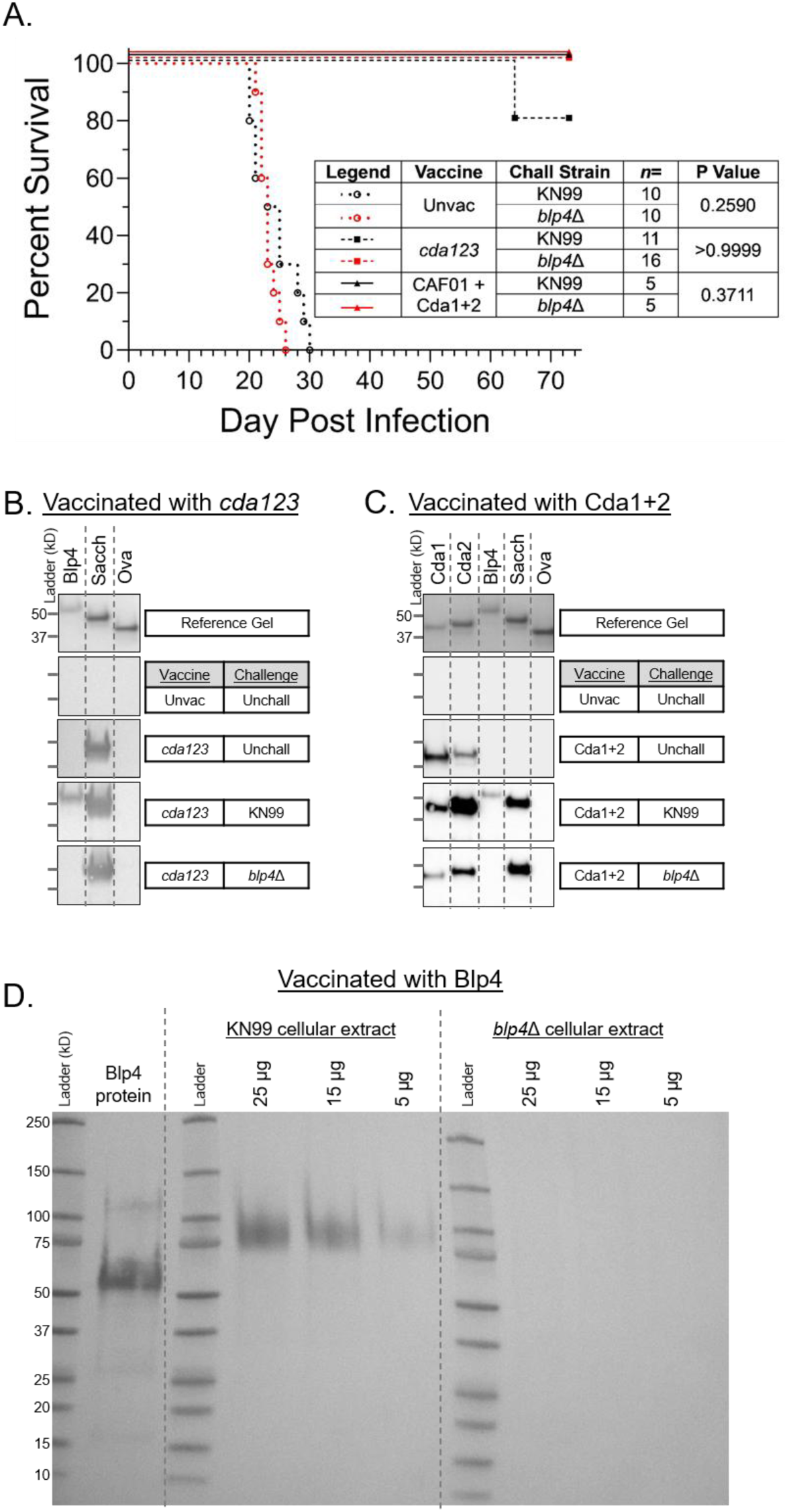
Specificity of the IgG antibody generated by vaccinated and infected mice for Blp4. BALB/c mice were either vaccinated with live *cda123* or Cda1 and Cda2 (Cda1+2) adjuvanted with CAF01 before receiving an orotracheal challenge of 2 x 10^4^ CFU of the indicated cryptococcal strain. (A) Survival was measured until 73 days post infection. Serum was collected from mice vaccinated with *cda123* (B) or Cda1+2 (C). Naïve (Unvac, Unchall) serum was used as a negative control. Vaccinated, unchallenged (Unchall) serum was collected two weeks after the final vaccination. Vaccinated mice were challenged orotracheally with either KN99 or *blp4*Δ, and serum was collected from mice surviving until 73 DPI. (D) BALB/c mice were vaccinated with Blp4 adjuvanted with CAF01. Serum was collected two weeks after the final vaccination and then blotted against either Blp4 recombinant protein or titrations of cryptococcal cellular extracts from KN99 and *blp4*Δ. Mouse serum was used as the primary antibody with either alkaline phosphatase-conjugated α-IgG (B and D) or HRP-conjugated α-IgG (C) as a secondary antibody.

Blp4 antibody specificity was first probed using the sera of mice vaccinated with live *cda123* and left unchallenged, or vaccinated and then challenged with either KN99 or *blp4*Δ. Antibody responses were then measured by Western blotting (**Figure 3B**). As in **Figure 2**, IgG antibody to Blp4 was absent in the serum of both naïve mice and vaccinated, unchallenged mice, but IgG to Blp4 was present in the serum of mice surviving KN99 challenge. However, mice challenged with *blp4*Δ did not generate an IgG response to Blp4 (**Figure 3B**). Also consistent with the results shown in **Figure 2**, all vaccinated mice developed an antibody response to Sacch but not to the recombinant Ova control. Results were similar when using a recombinant protein vaccine: Unchallenged mice vaccinated with Cda1 and Cda2 adjuvanted with CAF01 (Cda1+2) developed serum IgG only to the vaccine antigens Cda1 and Cda2, but not to Blp4. Subsequent challenge with KN99 generated antibody to Blp4 protein as seen with serum from *cda123* vaccinated mice, but again not with serum from mice challenged with the *blp4*Δ strain (**Figure 3C**).

To further investigate specificity of the antibody response to Blp4, we next compared cellular extracts from wild-type *C. neoformans* strain KN99 and the *blp4*Δ strain. We examined, by Western blotting, whether serum from mice immunized with Blp4 had IgG antibodies to recombinant Blp4 and cellular protein from the two extracts (**Figure 3D**). We found the immune serum reacted with the recombinant Blp4 and extracts from strain KN99, but there was no detectable antibody binding to proteins extracted from *blp4*Δ. The higher molecular size of the band seen with the KN99 cell extract is presumably due to posttranslational modification of Blp4, particularly glycosylation, which is found in fungal but not *E. coli*-derived proteins [27]. Thus, by Western blotting, Blp4 antibody does not cross-react with other members of the Blp family in *Cryptococcus*.

A major mechanism by which IgG antibody can contribute to protection against infection is by promoting opsonophagocytosis. Therefore, we next investigated whether serum from naïve and immunized mice promoted phagocytosis of mCherry-labeled KN99 by bone marrow-derived macrophages (BMDMs). Solophenyl, which does not penetrate into BMDMs [28], was used to identify yeast cells that were not internalized (**Figure 4A**). The serum was heat-inactivated to eliminate complement opsonization as a variable. Although the effects were somewhat modest, opsonization of *C. neoformans* with Blp4-vaccinated serum yielded a significantly higher phagocytic index than that seen with naïve serum and with serum from mice immunized with Ova, Cda1+2, and Sacch (**Figure 4B, 4C**).

**Figure 4:**
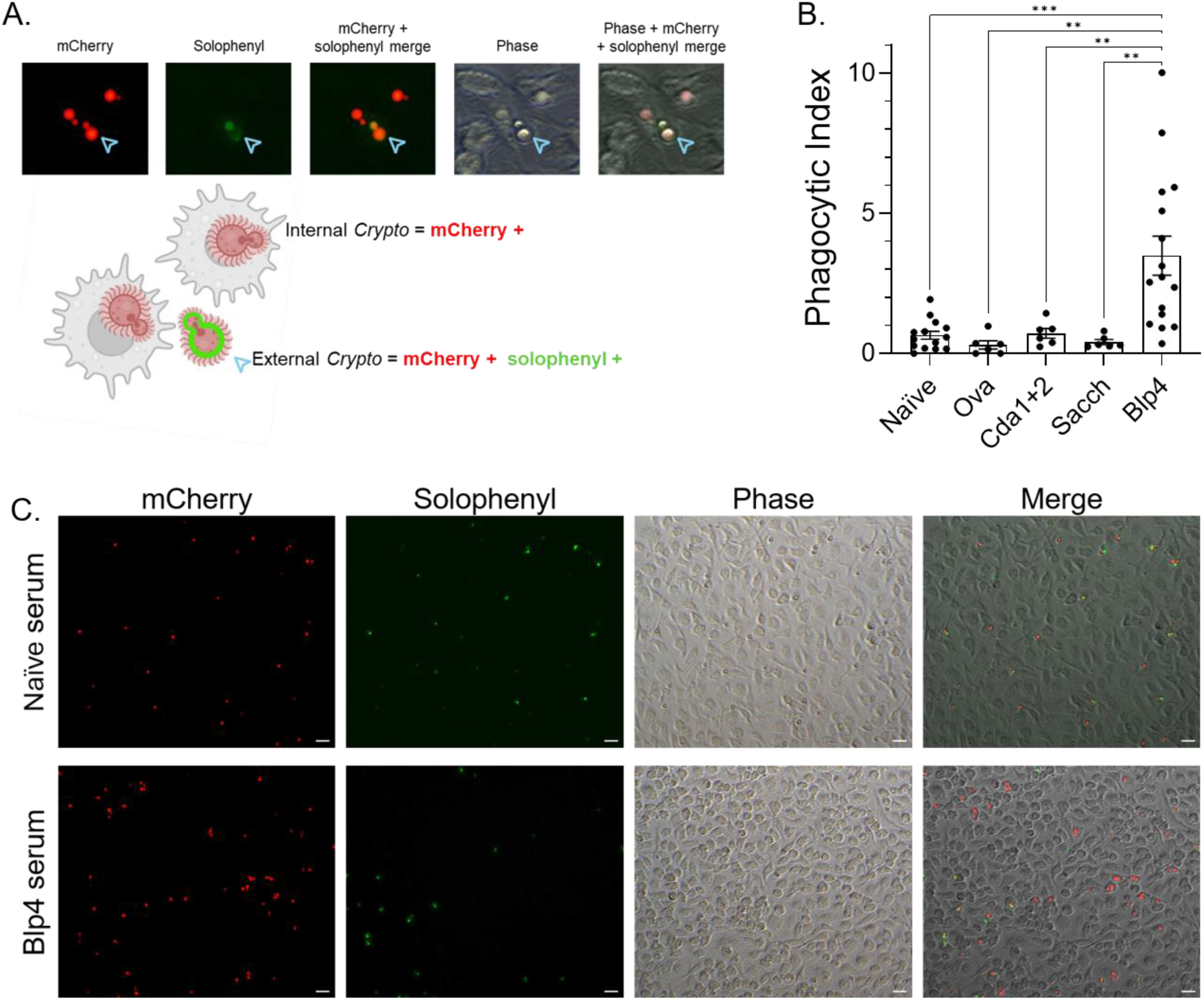
Effect of opsonization of *C. neoformans* with serum from Blp4-immunized mice on phagocytosis by murine macrophages. BMDMs were incubated for two hours with mCherry *C. neoformans* KN99 at a 1:1 ratio in the presence of 10% heat-inactivated mouse serum. Solophenyl was added for the last 15 minutes. Free fungi were washed away and phagocytosis assessed by fluorescent microscopy. (A) Schematic and representative images of the phagocytosis assay. Solophenyl stains extracellular but not phagocytosed (internalized) *C. neoformans.* (B) Phagocytic index (PI) was calculated as internal *C. neoformans* cells per 100 macrophages. Serum was from mice that were naïve or immunized with the antigens indicated on the x-axis. Each dot represents the PI of one infected well of macrophages. For each well, three random fields were quantified. Data are representative of 2-8 experiments, each containing at least two wells per condition. (C) Representative images taken using a 200X magnification. Scale bar is 20 μM.

The above studies utilized *C. neoformans* KN99, which is virulent reference strain derived from a clinical strain isolated in the United States from a patient with lymphoma [29]. We recently demonstrated that *cda123* vaccination protected mice from challenge with five African and Asian *C. neoformans* clinical strains (designated, BK80, BMD1338, PMH1063, PMH1065, and UgCl395) [8]. Using the serum from vaccinated BALB/c mice that survived challenge with these strains, we assayed the IgG antibody response to our panel of cryptococcal antigens by dot blot (**Figure 5**). Consistent with what we found with KN99 challenge, there was a robust IgG antibody response to Blp4 and Sacch in the serum of mice challenged with each of the five clinical strains (**Figure 5**). IgG antibody responses to other cryptococcal antigens were seen in serum from mice infected with some of the strains.

**Figure 5:**
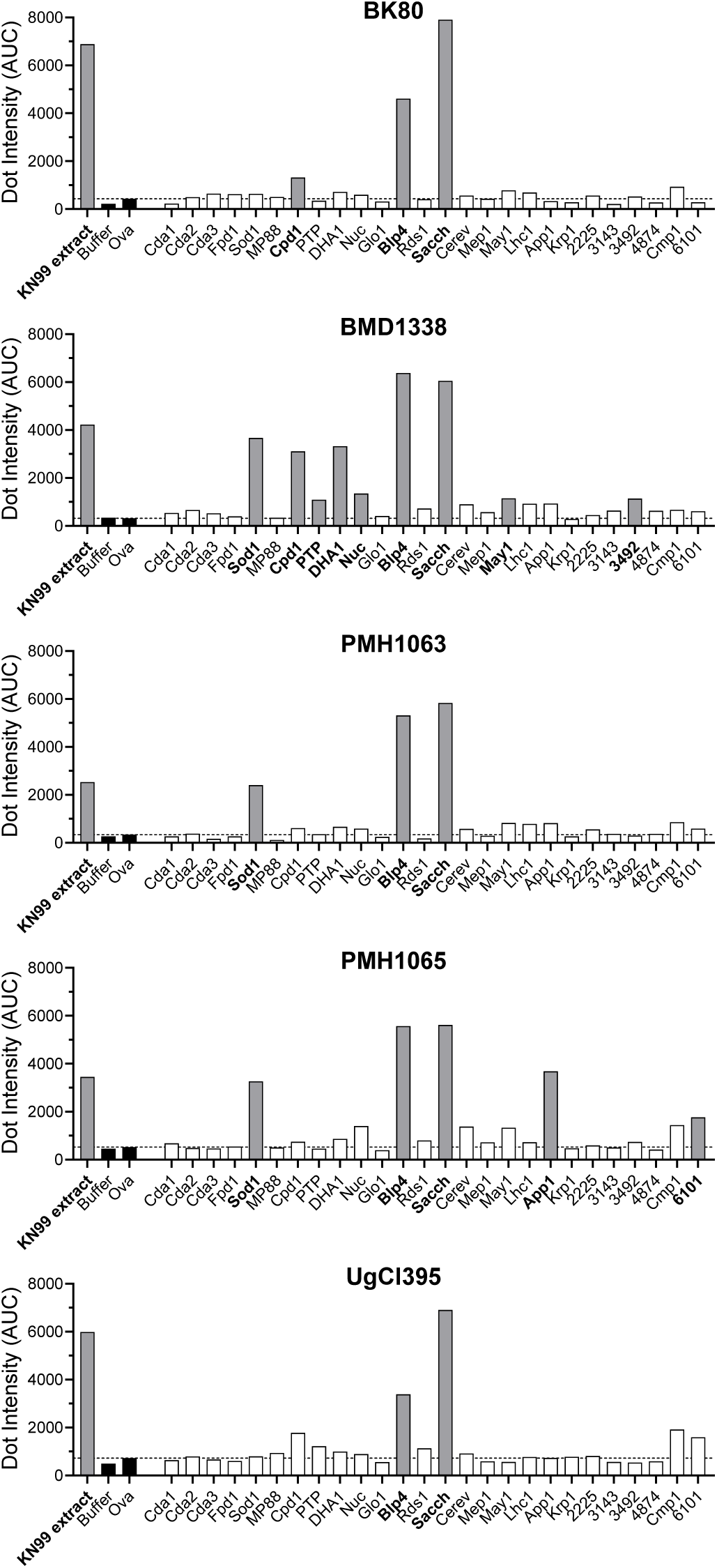
Antibody to cryptococcal antigens in *cda123*-vaccinated mice challenged with *C. neoformans* clinical strains. BALB/c mice were vaccinated with live *cda123* before challenge with the indicated clinical strain. Serum was collected from surviving mice at day 70 post infection and pooled by clinical strain (n=2-5 mice/group). Dot blots were done as in Figure 2. Mouse serum was used as the primary antibody, and alkaline phosphatase-conjugated IgG was used as the secondary antibody. Dot intensity was measured using ImageJ. Negative controls (buffer only, Buffer; and recombinant ovalbumin, Ova) are denoted by black bars. The dotted line indicates the dot intensity of the Ova control. Dot intensity 2.5X higher than that of Ova was considered positive and indicated by grey bars.

Lastly, we sought to determine whether patients with cryptococcosis also developed IgG antibodies against Blp4 (**Figure 6**). To do this, we took advantage of the availability of serum samples from subjects enrolled in an ongoing National Institutes of Health clinical study of patients who developed cryptococcosis in the absence of known immunocompromising conditions that would predispose them to cryptococcosis. Details of the study can be found at Study Details | NCT00001352 | Fungal Infection Susceptibility | ClinicalTrials.gov. Initial screening probed a pool of four patient sera against our panel of cryptococcal antigens. We previously reported on the peripheral blood mononuclear cell (PBMC) response to cryptococcal antigens for the individual donors in this pool [13]. We found that, consistent with the mouse data, the pooled patient sera reacted to Blp4 and Sacch. A third antigen, phosphotidylinositol transfer protein (PTP), was also identified as binding IgG in pooled patient sera. PTP was initially screened as a recombinant antigen due to its abundance in protective cryptococcal alkaline extract vaccines [10, 30]; however, we were unable to demonstrate protection in either BALB/c or C57BL/6 mice vaccinated with GP-PTP [10].

**Figure 6:**
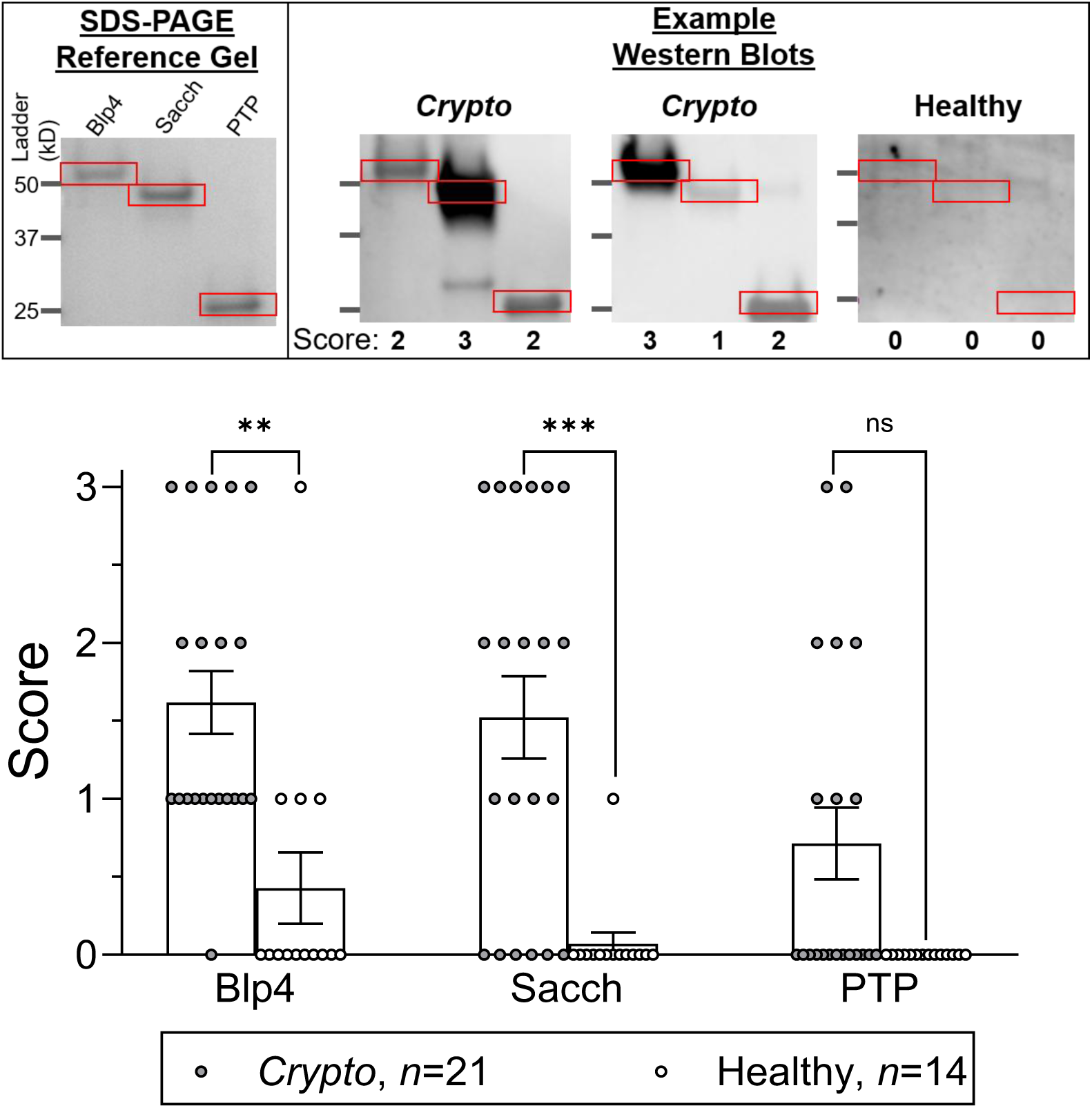
Antibody to cryptococcal antigens in human sera. Individual human sera samples were probed via Western blot for antibody to Blp4, Sacch, and PTP. Protein size was visualized via SDS-PAGE (top left). Band intensity was quantified for each antigen on immunoblots from both patients with a history of cryptococcosis, and healthy control donors, by blind scoring by three reviewers. A score of 0 is negative, while 3 is the most positive. Example Western blots and the corresponding scores for each antigen are shown above. The red boxes indicate the correct molecular weight for the recombinant protein. The median score for the 3 reviewers was graphed. Individual blots and scores are shown in **Supplementary Figure 6** and **Supplementary Figure 7**. Each dot represents the median score for an individual serum donor. Statistical significance was determined by unpaired t tests with the Bonferroni correction. ns, not significant at P>0.05; ** significant at P≤0.01; *** significant at P≤0.001.

We then probed the IgG response to Blp4, Sacch, and PTP in individual serum samples obtained from 21 subjects with cryptococcosis and 14 healthy subjects (**Figure 6**). Details regarding the serum donors, including demographic, clinical, and immunologic information on the subjects with cryptococcosis, can be found in **Supplementary Table 1**. We established a qualitative scoring system based on the intensity of the band on the Western blot, with 0 being negative and 3 being the most positive. Blots were blindly scored by three individual reviewers, and the median of the three assigned scores was graphed. Examples of blots and scores are shown at the top of **Figure 6**. Individual blots for all samples, with the median scores, are shown in **Supplementary Figure 6 and Supplementary Figure 7**. Twenty of the 21 (95.2%) patient sera tested had IgG to Blp4 compared with only four of the 14 (28.6%) healthy sera (p<0.001). For Sacch, the comparison between patient and control sera was also highly significant, with 15/21 (71.4%) and 1/14 (7.1%) sera, respectively, containing detectable IgG to Sacch (p<0.0001). Finally, 8/21 (38.1%) patient sera contained IgG that bound PTP compared with 0/14 (0%) control sera; however, the difference between groups was not statistically significant. Thus, in addition to vaccinated mice with experimental cryptococcosis having IgG responses to Blp4, human subjects who survived cryptococcosis consistently do as well.

## Discussion

As most persons at risk for cryptococcosis have impaired T cell function, successful vaccines will likely need to elicit significant antibody-mediated protection. The carbohydrate capsule of *Cryptococcus* is an attractive target; however, shed GXM, which has a very long half-life, can form complexes with antibody, thereby competing with GXM on the yeast cell [20]. Recent studies identified four cryptococcal proteins, Blp4, Cpd1, Lhc1, and Krp1 as being capsule-associated [21, 25]. We had already demonstrated that vaccines containing Blp4, Cpd1, and Lhc1 protect wild-type mice [10, 11, 14]. Here, we extended this observation by showing that Krp1 is also a protective antigen. To test the hypothesis that antibodies contributed to protection, we compared mortality curves in wild-type and B cell-deficient mice vaccinated with Blp4, Cpd1, Lhc1, and Krp1 and then infected with *C. neoformans*. We found that protection is diminished in mice lacking B cells. In contrast, there was no mortality phenotype in B cell-deficient mice vaccinated with Cda1 or Cda2 (**Figure 1** and [12]), two antigens that are present in extracellular vesicles [24] and not known to be specifically capsule-associated. As with Blp4, Cpd1, Lhc1, and Krp1, vaccination with Cda1 and Cda2 adjuvanted in CAF01 elicited an antibody response, albeit one that apparently did not significantly contribute to protection.

Antibody to Blp4 was observed in the serum of mice vaccinated with *cda123* and infected with *C. neoformans*. The antibody response was found in two strains of mice and following challenge with six *C. neoformans* strains, including five clinical strains isolated from patients in Africa and Asia. Blp4 is part of a family of six cryptococcal proteins containing a Barwin-like domain. However, the antibody was specific to Blp4 as no antibody response to Blp4 was observed when vaccinated mice were infected with a *C. neoformans* strain deleted only of the *BLP4* gene.

In contrast, a serum antibody response to Cpd1, Lhc1, and Krp1 was not seen in *cda123*-vaccinated mice challenged with *C. neoformans* strain KN99. However, antibodies to Cpd1 were seen following infection with two of the clinical strains tested. Interestingly, in an immunoblot screen of 26 recombinant proteins, most of which we have tested as vaccine antigen candidates, anti-Sacch antibodies were found in *cda123*-vaccinated mice, even prior to KN99 challenge. Despite Sacch being a prevalent IgG antibody component in mouse serum, vaccines containing Sacch adjuvanted in GPs are not protective in C57BL/6 mice [10]. The lack of protection may be due in part to Sacch not being surface-exposed, as we demonstrated that serum containing anti-Sacch antibodies does not promote opsonophagocytosis.

Blp4 has several attractive features which led us to include it as part of a published panel of cryptococcal vaccine antigens [10]. First, it has a serine/threonine-rich region which is a presumed site of extensive O-linked mannosylation [31]. Second, Blp4 is present in the cryptococcal secretome and transcriptome, evidenced by RNA-seq analysis of *Cryptococcus* isolated from the CSF of human patients [32-34]. Third, Blp4 shares no human homology [10], which should reduce the risk of autoimmune reactions following vaccination. Fourth, Blp4 shares a conserved double psi beta barrel domain with five other cryptococcal antigens and belongs to the Barwin-like protein family in *Cryptococcus*, comprised of Blp1 through Blp6 [26]. We have prioritized selection of candidate antigens belonging to protein families to promote cross-reactivity to additional cryptococcal antigens [10, 11], however we show here that Blp4 does not appear to be cross-reactive. Other features that make Blp4 a promising vaccine candidate include its identification in cryptococcal extracellular vesicles [24] and, as noted above, its association with the outer portion of the capsule [21].

Our published work demonstrating protection of BALB/c and C57BL/6 mice by Blp4 delivered in glucan particle vaccines [10] was extended in the present study by showing that vaccines containing Blp4 adjuvanted with CAF01 were also protective. CAF01, a liposome-based adjuvant containing the glycolipid Mincle agonist α,α′-trehalose 6,6′-dibehenate, promotes adaptive Th1, Th17, and antibody responses to vaccine antigens [14, 35]. This has translational relevance because CAF01 has a record of safety and efficacy in human clinical trials [36, 37].

Other groups have screened mouse, human, and koala serum samples for specific cryptococcal antibodies without identifying antibodies bound to Blp4 [38-44]. We screened serum samples against a panel of recombinant *Cryptococcus* proteins. While this approach is limited to the proteins being screened, it should have higher sensitivity to detect an antibody response to antigens that are included in the screen. In addition, secreted cryptococcal proteins are often highly glycosylated [45]. Such glycoproteins, especially those that, like Blp4, have serine/threonine rich regions, may not run as distinct spots following two-dimensional gel immunoelectrophoresis [46]. Glycosylation also can mask peptide epitopes.

Heat-inactivated serum obtained from mice immunized with Blp4 promoted macrophage phagocytosis of *C. neoformans*. Serum antibodies to Sacch were detected following mouse vaccination with live *cda123*, in mice challenged with KN99, and in humans with cryptococcosis. However, serum from mice immunized with Sacch did not promote opsonophagocytosis of *Cryptococcus*. Similarly, sera from mice vaccinated with Cda1 and Cda2 did not promote opsonophagocytosis. Although vaccination with formulations containing Cda1 and Cda2 induce the production of specific antibody, protection is CD4^+^ T cell-mediated [11, 12, 14]. Taken together, these data emphasize the importance of location on the capsular surface for opsonophagocytosis, and not antigenicity alone, in antibody cell-mediated protection. Phagocytosis could promote host defenses by leading to antimicrobial activity and enhancing T cell responses via augmentation of MHC Class-II antigen presentation [47-49].

The clinical significance of finding serum antibodies to Blp4, Sacch, and PTP in subjects with a history of cryptococcosis merits further study. The study subjects who donated serum differed from most patients with cryptococcosis in that an exclusion criterion for entry into the study was known immunodeficiency. It is unknown whether serum from immunocompromised or immunosuppressed patients with cryptococcosis, such as persons living with HIV and recipients of solid organ transplants, would have similar antibody responses and may be worth exploring in further studies. Similarly, it would be worthwhile to study whether the antibody responses to Blp4, Sacch, and PTP have diagnostic or prognostic significance.

Recent work from our lab demonstrated that following incubation with Blp4, human PBMCs isolated from cryptococcosis patients have significantly higher CD4^+^ T cell stimulation indices compared with healthy controls [13]. Moreover, we have mapped stimulatory CD4^+^ T cell epitopes in the vaccine candidate antigens Cda1 and Cda2 and created a multiepitope vaccine that protects mice expressing humanized MHC Class II molecules [50]. Ongoing studies in our lab are focused on the identification of CD4^+^ T cell and B cell epitopes in Blp4 with plans to include these epitopes in multiepitope vaccines designed to elicit both CD4^+^ T cell and antibody mediated protection. The majority of patients with cryptococcal infections have impaired CD4^+^ T cell function. Primary B cell immunodeficiency as the primary risk factor for cryptococcosis is rare, although perturbated humoral immunity may be seen [18, 51]. These findings suggest that effective vaccination against *Cryptococcus* may be achievable by targeting immune pathways that remain functional in immunocompromised hosts. Accordingly, a key strategy is to vaccinate at risk populations, such as people living with HIV and those on solid organ transplant lists, while CD4⁺ T cell levels are still relatively preserved. This would allow for durable protective immunity to be established through complementary CD4^+^ T cell and antibody effector mechanisms. If CD4^+^ T cell function declined, protection could be maintained via an antibody response specific to Blp4. Taken together, the findings that Blp4:1) stimulates strong antibody and CD4^+^ T cell responses in mice and humans with cryptococcosis, 2) is a protective antigen when formulated in preclinical subunit vaccines, 3) is abundantly transcribed during infection, 4) is associated with the cryptococcal capsule, and 5) has no human homology, lend strong support for its inclusion in subunit cryptococcal vaccines.

## Materials and Methods

### Chemicals and culture media

Reagents were from Thermo Fisher Scientific (Pittsburgh, PA) unless otherwise stated. *Cryptococcus* culture media was YPD (Difco yeast extract, Bacto peptone, Dextrose, with and without 2% agar), and Sabouraud Dextrose agar. The BL21 strain of *E. coli* was used for protein expression and cultured for 18 hours at 30°C with shaking in Overnight Express Instant TB medium (MilliporeSigma, Burlington, MA), as in prior studies [9-12]. Ampicillin at 100 μg/mL, was used for antibiotic selection.

### Strains of Cryptococcus

*C. neoformans* strains (**Table 1**) were maintained as glycerol stocks at -80°C. Initial culture plates were grown on YPD agar and used to inoculate liquid YPD medium. For *cda123* vaccination, strains were prepared as previously described [7, 52, 53]. Briefly, 20 mL and 4 mL liquid YPD cultures for orotracheal (OT) vaccination and subcutaneous boosts, respectively, were inoculated and shaken at 225 RPM for two days at 30°C. Strains used for fungal challenges were cultured in 4 mL liquid YPD media with shaking for 18 hours. Cells were harvested via centrifugation, washed with PBS, and counted using an automated cell counter (BioRad TC20, Hercules, CA). Concentrations were adjusted in PBS to 2 x 10^8^ cells/mL for *cda123* OT vaccination, 2 x 10^7^ cells/mL for *cda123* subcutaneous vaccination, 4x10^5^ cells/mL for OT fungal challenge of mice on the BALB/c background, and 2 x 10^5^ cells/mL for mice on the C57BL/6 background. All OT fungal vaccinations and challenges were verified by plating on Sabouraud dextrose agar.

**Table 1:**
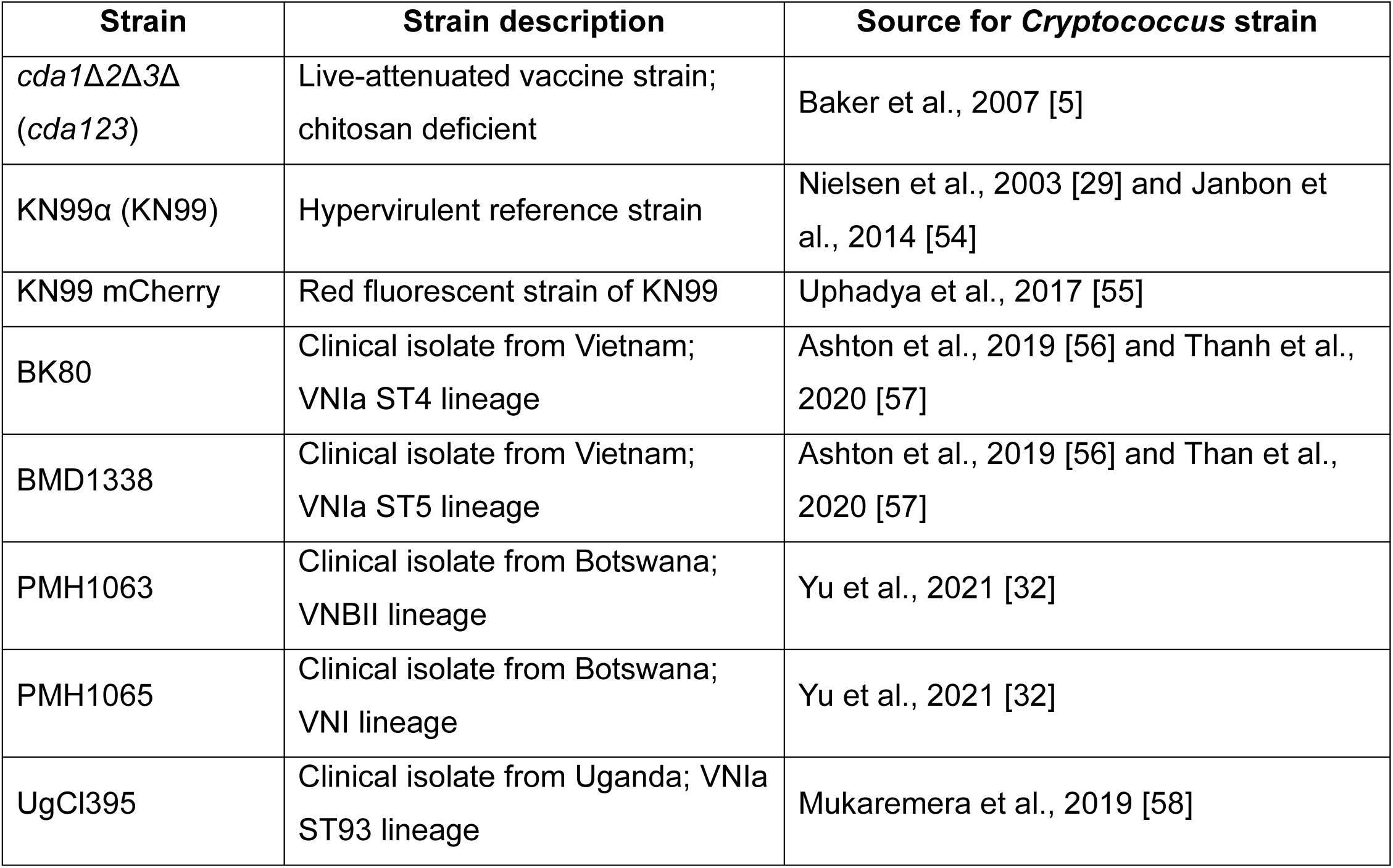
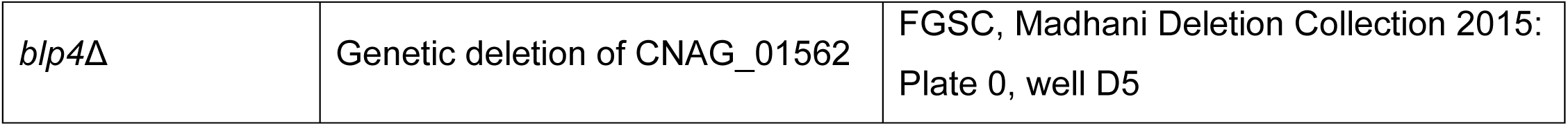
*C. neoformans* strains used.

| Strain | Strain description | Source for *Cryptococcus* strain |
|---|---|---|
| *cda1*Δ*2*Δ*3*Δ (*cda123*) | Live-attenuated vaccine strain; chitosan deficient | Baker et al., 2007 [5] |
| KN99α (KN99) | Hypervirulent reference strain | Nielsen et al., 2003 [29] and Janbon et al., 2014 [54] |
| KN99 mCherry | Red fluorescent strain of KN99 | Uphadya et al., 2017 [55] |
| BK80 | Clinical isolate from Vietnam; VNIa ST4 lineage | Ashton et al., 2019 [56] and Thanh et al., 2020 [57] |
| BMD1338 | Clinical isolate from Vietnam; VNIa ST5 lineage | Ashton et al., 2019 [56] and Than et al., 2020 [57] |
| PMH1063 | Clinical isolate from Botswana; VNBII lineage | Yu et al., 2021 [32] |
| PMH1065 | Clinical isolate from Botswana; VNI lineage | Yu et al., 2021 [32] |
| UgCl395 | Clinical isolate from Uganda; VNIa ST93 lineage | Mukaremera et al., 2019 [58] |
| *blp4*Δ | Genetic deletion of CNAG_01562 | FGSC, Madhani Deletion Collection 2015: Plate 0, well D5 |

### Expression of cryptococcal proteins in *E. coli* and protein purification

App1, Krp1, 2225, Cmp1, and 6101 are proteins previously unpublished in our studies and were synthesized and cloned in pET19 by Genscript (Piscatway, NJ, USA). All recombinant proteins were expressed in *E. coli* (Table 2) and purified on His-Bind resin (MilliporeSigma) as previously described [10]. Briefly, BL21 *E. coli* (New England Bio Labs, Ipswich, MA) was transformed with a plasmid expressing the selected recombinant protein and then cultured in Overnight Express Instant TB medium (MilliporeSigma). Cells were lysed via a series of freeze-thaw cycles and sonication. Protein was solubilized in 6M urea and purified over a His-Bind nickel column. All purification buffers contained 6M urea to maintain solubility of the proteins. Following SDS-PAGE analysis, eluted fractions containing the proteins of interest were pooled, dialyzed to remove imidazole, and concentrated to either 10 mg/mL or 5 mg/mL. Protein stocks were stored at -80°C until use.

**Table 2:** Cryptococcal antigens used.

| Antigen | CNAG | Description | References/Citations |
|---|---|---|---|
| Cda1 | CNAG_05799 | Chitin deacetylase 1 | [5, 9, 10] |
| Cda2 | CNAG_01230 | Chitin deacetylase 2 | [5, 9, 10, 30] |
| Cda3 | CNAG_01239 | Chitin deacetylase 3 | [5, 9, 10] |
| Fpd1 | CNAG_06291 | Fungal polysaccharide deacetylase 1 | [5, 9, 10] |
| Sod1 | CNAG_01019 | Superoxide dismutase 1 | [9, 10, 30] |
| MP88 | CNAG_00776 | Mannoprotein 88 | [9, 10] |
| Cpd1 | CNAG_00919 | Carboxypeptidase 1 | [10, 11, 21, 30] |
| PTP | CNAG_07442 | Phosphotidylinositol transfer protein | [10, 30] |
| DHA1 | CNAG_07422 | Delayed-type hypersensitivity protein 1 | [10, 59] |
| Nuc | CNAG_00264 | S1 nuclease/phospholipase C | [10, 30] |
| Glo1 | CNAG_02030 | Glyoxal oxidase 1 | [10] |
| Blp4 | CNAG_01562 | Barwin-like protein 4 | [10, 21] |
| Rds1 | CNAG_06267 | Rds1-like protein | [10] |
| Sacch | CNAG_00581 | Saccharopepsin | [10, 30] |
| Cerev | CNAG_04625 | Cerevisin | [10, 30] |
| Mep1 | CNAG_04735 | Metallo proteinase 1 | [10, 11] |
| May1 | CNAG_05872 | Major aspartyl peptidase 1 | [10, 60] |
| Lhc1 | CNAG_04753 | Lactonohydrolase 1 | [10, 11, 21] |
| App1 | CNAG_06574 | Antiphagocytic protein 1 | [61] |
| Krp1 | CNAG_05595 | Kelch repeat-containing protein 1 | [25] |
| 2225 | CNAG_02225 | Glucan 1,3-beta-glucosidase | [62] |
| 3143 | CNAG_03143 | Heat shock protein 9/12 | [10] |
| 3492 | CNAG_03492 | Unknown | [10] |
| 4874 | CNAG_04874 | Glucanase | [10] |
| Cmp1 | CNAG_06000 | Cryptococcus mannoprotein 1 | [32] |
| 6101 | CNAG_06101 | ADP/ATP carrier protein | [63] |

### Cryptococcal whole cell protein extracts

From stock plates of either KN99 or *blp4*Δ, 4 mL tubes of liquid YPD were inoculated and cultured overnight at 30°C with shaking. 100 μL were subcultured into 20 mL of liquid YPD media in a 250 mL vented flask and cultured under the same conditions for an additional 48 hours. The cultures were then divided into 2 mL screw-cap tubes and processed on ice. The cells were collected via centrifugation, and the supernatant was discarded. Cells were resuspended in 1 mL of Roche cOmplete Protease Inhibitor Cocktail with 0.05% Triton-x100. The equivalent volume of 0.6 mL of 0.7mm Zirconia beads was added to each tube followed by two rounds of bead beating for 2.5 minutes with a MiniBeadBeater-16, Model 607 (BioSpec Products, Bartlesville, OK). Cell lysis was confirmed via microscopy before proceeding. The liquid portion of the cell slurry was gently removed from the beads and centrifuged at 20,800 x g to collect the supernatant. The protein concentration of the extracts was determined with bicinchoninic acid using the Pierce BCA Protein Assay Kit, and the protein extracts were stored at -80°C.

### Vaccination and challenge

All subcutaneous vaccinations were delivered as 0.1 mL doses in the abdomen. For GP-vaccines, mice received a series of three vaccinations, with two weeks between vaccinations. GP-vaccines were prepared as previously described [9, 10]. Each vaccination with CAF01 was delivered in two 100 μL doses, with one injection into each side of the abdomen. Each dose contained 5 μg of protein and CAF01 (Statens Serum Institut, Copenhagen, Denmark) [14, 50]. The second vaccination followed three weeks later and was delivered in the same manner. Live vaccinations with *cda123* were prepared as previously described and delivered as one orotracheal vaccination of 1 x 10^7^ cells, followed by two subcutaneous vaccinations of 2 x 10^6^ cells, each two weeks apart [6-8, 53]. Two weeks after the final immunization, mice were challenged with the indicated cryptococcal strain. Mice on the C57BL/6 background received an orotracheal challenge dose of 1 x 10^4^ cells, and BALB/c mice received a challenge dose of 2 x 10^4^ cells [10]. Mice were monitored daily for survival.

### Mice

BALB/c (Strain #000651), C57BL/6 (Strain #000664), and muMT (Strain #002288) mice were purchased from the Jackson Laboratory (Bar Harbor, ME). Male and female mice were used in approximately equal numbers for all experiments. Mice were housed and bred in a specific pathogen-free environment at the University of Massachusetts Chan Medical School (UMCMS). All experimental protocols were approved by the UMCMS Institutional Use and Care of Animals Committee.

### Human samples

Human blood from subjects with cryptococcosis and healthy controls was obtained by venipuncture under experimental protocols approved by the Institutional Review Boards of the National Institutes of Health and UMCMS. Sera were collected following centrifugation of clotted blood and stored at -80°C.

### Immunoblots

Mouse blood was collected at the indicated time points by cardiac puncture of fully anesthetized mice. For Western blots, recombinant protein was electrophoresed into a 4-20% gradient gel (BioRad) in Tris-glycine-SDS running buffer. Precision Plus Dual Color Standards (BioRad) were used to estimate molecular weight on SDS-PAGE gels. All reference gels contained 1 μg of protein per lane and were stained with Coomassie InstantBlue (Abcam, Cambridge, United Kingdom). For all Western blots, replicate gels containing the protein amounts indicated in **Table 3** were blotted onto nitrocellulose membranes (BioRad) using a Trans-Blot Turbo Transfer System (BioRad). For dot blots, protein was applied to a nitrocellulose membrane using the Bio-Dot Microfiltration Apparatus (BioRad). Protein samples were filtered through the membrane using gravity filtration; each well was then washed thoroughly with Tris-buffered saline with 1% Tween 20 (TBST) and removed from the apparatus. All blots were blocked overnight in TBST containing 5% non-fat dry milk at 4°C before incubation with the indicated pooled serum at a dilution of 1:500. The indicated secondary antibody from **Table 3** was applied at a 1:5,000 dilution before development with either SuperSignal West Pico PLUS Chemiluminescent Substrate or 1-Step NBT/BCIP for chemiluminescent or alkaline phosphatase detection, respectively. Chemiluminescent blots were imaged using the ChemiDoc Imaging System (BioRad), and alkaline phosphatase blots were imaged using the Gel Doc XR System (BioRad). For dot blots, dot intensity was quantified using ImageJ 1.54i [64].

**Table 3:** Western Blot Information.

| **Blot method** | **Detection method** | **Serum source** | **Recombinant protein/well** | **Secondary antibody** |
|---|---|---|---|---|
| Western blot | Chemiluminescence | Mouse | 0.1 µg | Goat anti-mouse IgG (H+L) secondary antibody, HRP (Invitrogen 31430) |
|  |  | Human | 0.5 µg | Goat anti-human IgG (H+L) secondary antibody, HRP (Invitrogen 31410) |
|  | Alkaline phosphatase | Mouse | 1 µg | Goat anti-mouse IgG, H&L chain Alkaline Phosphatase Conjugate (Calbiochem 401212) |
| Dot blot | Alkaline phosphatase | Mouse | 2 µg | Goat anti-mouse IgG, H&L chain Alkaline Phosphatase Conjugate (Calbiochem 401212) |

### Opsonophagocytosis assay

Bone marrow derived macrophages (BMDMs) were generated for opsonophagocytosis assays. Bone marrow cells were collected from mouse femurs and cultured with R10 media (RPMI supplemented with 10% fetal bovine serum, 1% GlutaMax, 0.55 mM Β-mercaptoethanol, 100 U/mL penicillin, and 100 μg/mL streptomycin) + 20 ng/mL M-CSF (GeminiBio, West Sacramento, CA) for 7-8 days, with media changes every three days. The BMDMs were then detached and seeded into flat-bottomed 96-well plates at a concentration of 10^5^ cells per well in 200 μL. The cells were allowed to attach overnight before non-adherent cells were gently washed off, and 10^5^ cells mCherry KN99 [55] plus 10% heat-inactivated (56°C, 30 minutes) mouse serum were added to each well. The assay proceeded at 37°C for one hour and 45 minutes before Solophenyl Flavine 7GFE 500 (Ciba Specialty Chemicals, Newport, DE), hereafter referred to as solophenyl, was added to each well at a final concentration of 10 μg/mL. After 15 minutes, free fungi were washed away, the remaining cells were fixed in 4% PFA, and imaging was done on a Nikon Eclipse TE200 microscope equipped with a SPOT InSight camera (SPOT Imaging, Serling Heights, MI). Each condition was run in duplicate. For each well, three random fields were imaged for quantification of phagocytosed KN99. Positive solophenyl staining was used to identify extracellular fungi. Phagocytic index was calculated as the number of internalized cryptococci per 100 macrophages.

### Statistics

Graphics and statistical analysis were conducted using GraphPad Prism V 10.4.2 (GraphPad Software, Boston, MA).

## Abbreviations

Blp: Barwin-like protein
BMDMs: bone marrow-derived macrophages
Cpd1: Carboxypeptidase 1
CAF01: Cationic Adjuvant Formulation 01
*cda123*: *cda1*Δ*cda2*Δ*cda3*Δ
Cda: chitin deacetylase
CFU: colony forming unit
CM: cryptococcal meningitis
DPI: day post infection
GP: glucan particle
GXM: glucuronoxylomannan
IFNγ: interferon gamma
Krp1: Kelch repeat-containing protein 1
Lhc1: Lactonohydrolase 1
Ova: Ovalbumin
PTP: Phosphotidylinositol transfer protein
Sacch: Saccharopepsin
Th1: T helper type 1

## Acknowledgements

The authors thank Zaida Ramirez-Ortiz for supplying some of the control human serum samples used in the study and Gabriel Kristian Pedersen (Statens Serum Institute) for supplying CAF01.

This work was supported, in part, by grants R24AI192252 (SML), R01AI172154 (SML), and R01AI125045 (SML, CAS), and contract 75N93019C00064, all awarded by the National Institute of Allergy and Infectious Diseases and by the Division of Intramural Research, NIAID, NIH Intramural Research Program, AI001124 (PRW) and AI001125 (PRW). The funders had no role in study design, data collection and interpretation, or the decision to submit the work for publication. The content of this publication does not necessarily reflect the views or policies of the Department of Health and Human Services, nor does mention of trade names, commercial products, or organizations imply endorsement by the US Government. The generation by the Madhani Laboratory and distribution of the deletion strains at the Fungal Genetics Stock Center was supported by NIH funding (R01AI100272).

