## Supplementary Figures and Table for "Identification of *Cryptococcus neoformans* capsule-associated protein antigens that mediate B cell-dependent vaccine protection in mice and elicit antibody responses in humans with cryptococcosis"

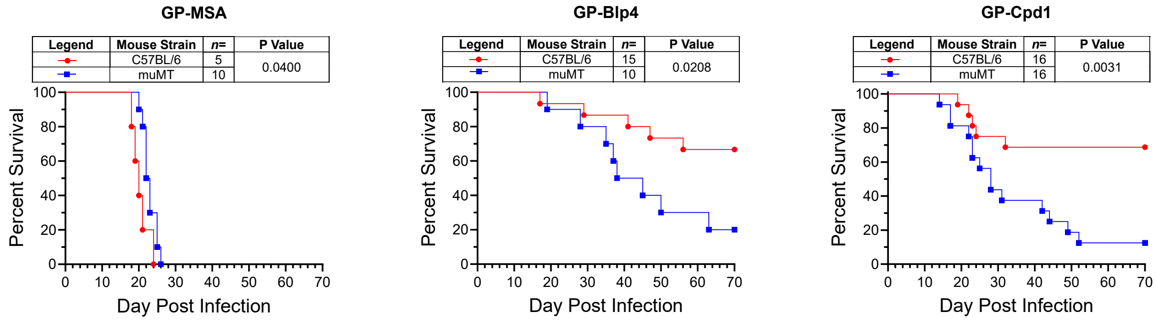


**Supplementary Figure 1: Contribution of B cells to protection by GP-vaccines.** Wild-type C57BL/6 mice and B cell-deficient (muMT) mice on the C57BL/6 background received a prime and two biweekly boosts with the indicated subcutaneous GP-based vaccine. GPs loaded with mouse serum albumin (MSA) were used as controls. Two weeks after the last boost, mice were challenged orotracheally with 1 x $10^4$ CFU of KN99 and followed for 70 days for survival. Kaplan-Meier survival curves were compared using the Mantel-Cox, log-rank test.

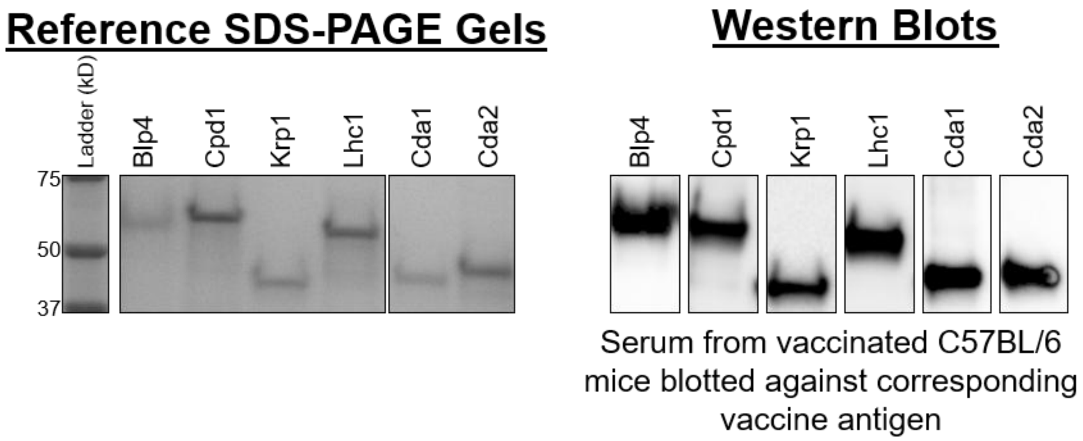


**Supplementary Figure 2: Western blots demonstrating specificity of IgG response to select vaccine antigens**. Mice were vaccinated as in **Figure 1**, and serum was collected from the vaccinated, unchallenged mice (day 0) two weeks after the final vaccination. The reference gel (left) shows the SDS-PAGE of the purified recombinant protein used in each CAF01 vaccine. Each lane of the reference gel contains 1 µg of recombinant protein stained with Coomassie, while each lane for the Western blots (right) contains 0.1 µg of recombinant protein. The label atop each blot indicates the recombinant protein used both for vaccination and the corresponding blot. Immune serum was used as the primary antibody, with HRP-conjugated α-mouse IgG as the secondary. Blots were developed with chemiluminescent substrate before imaging. Serum was pooled from groups of n=3-6 mice for each vaccination group. The Western blots have been cropped to show the relevant portions but have not been digitally manipulated.

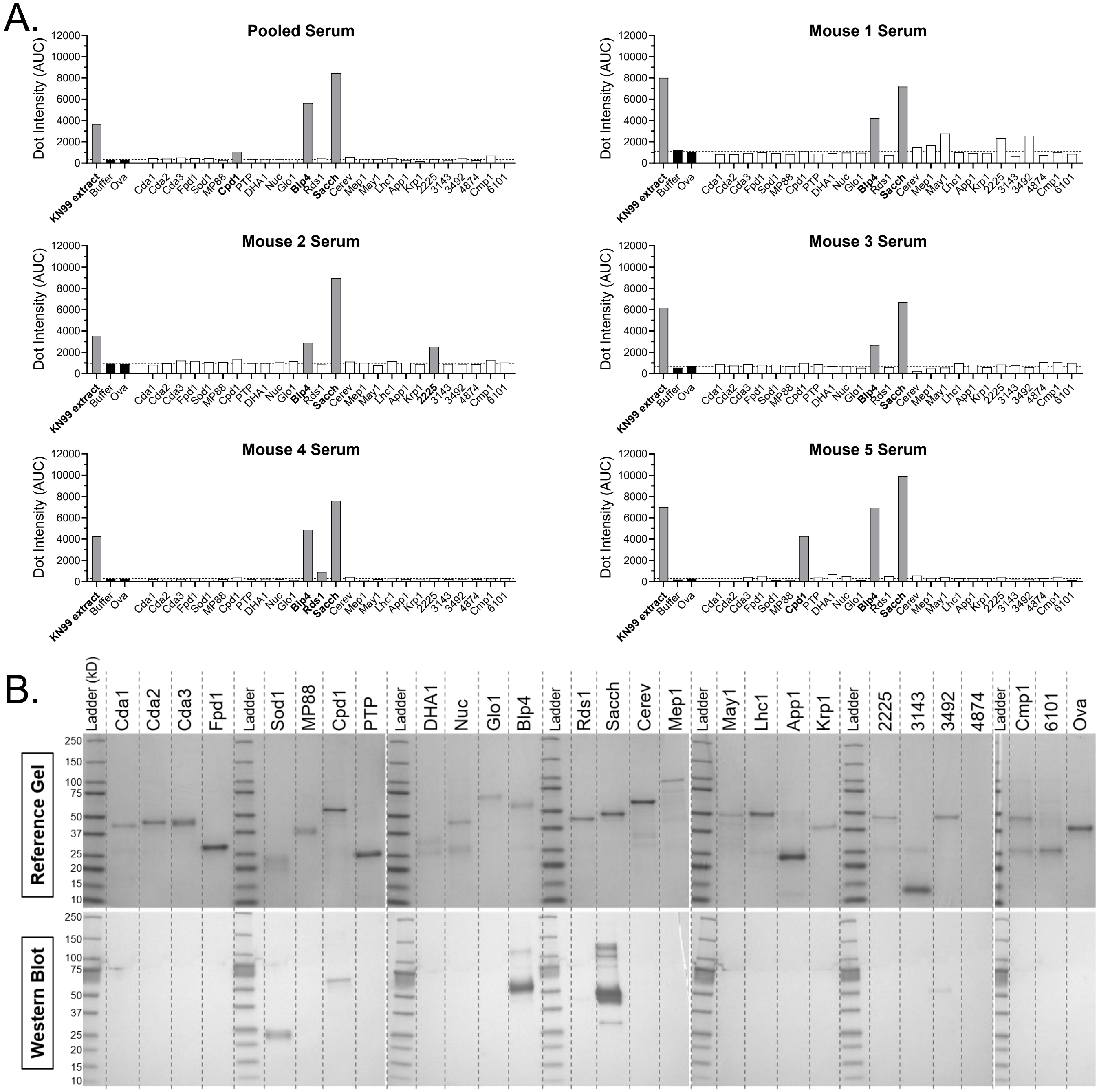


**Supplementary Figure 3: Dot blots as a semi-quantitative method for screening for the presence of antibody to cryptococcal antigens in vaccinated mice. (A)** Individual recombinant protein (2 µg), and extract of KN99 cellular protein (20 µg, grown in YPD), were spotted on nitrocellulose. Each blot was incubated with the pooled serum of Mouse 1 through Mouse 5, or the serum from each individual mouse, as a primary antibody; the five BALB/c mice were vaccinated with *cda123* and survived 70 days following KN99 challenge. Alkaline phosphatase-conjugated IgG was used as a secondary antibody. Dot intensity was measured using ImageJ, and the resulting area under the curve (AUC) was graphed. Negative controls (buffer only, Buffer; and recombinant ovalbumin, Ova) are denoted by black bars. The dotted line indicates the dot intensity of the Ova control. Dot intensity 2.5X higher than that of Ova were considered positive and indicated by grey bars. (B) The proteins screened via dot blot were analyzed by SDS-PAGE and Western blot. Top: The reference gel shows the SDS-PAGE gel of the recombinant proteins screened. Each lane contains 1 µg of protein, and the gel was stained with Coomassie. Bottom: Western blots used the pooled serum. Protein ladder is shown after every four lanes of recombinant protein. “Pooled serum” represents a pool of serum from Mouse 1 through Mouse 5.

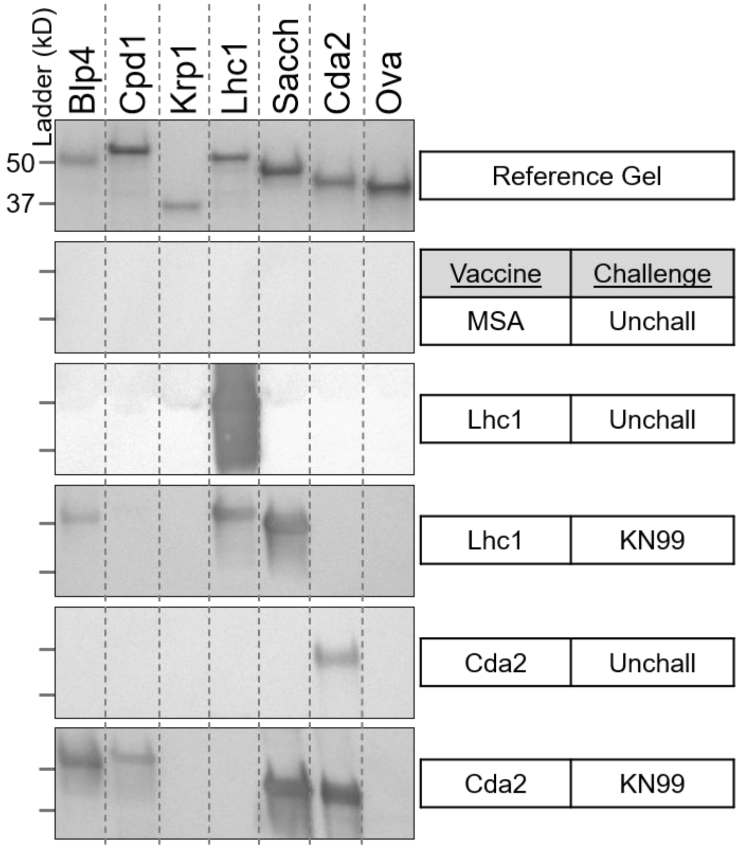


**Supplementary Figure 4: IgG antibody response against select cryptococcal antigens in GP-vaccinated mice challenged with KN99**. BALB/c mice were vaccinated with GPs containing MSA (control), Lhc1, or Cda2, as in **Supplementary Figure 1**. For unchallenged mice (Unchall), serum was collected 2 weeks after the final vaccination. For mice surviving KN99 challenge, serum was collected 70 days following infection. The reference gel shows the SDS-PAGE of Coomassie-stained purified cryptococcal recombinant proteins. Each lane contains 1 µg of protein. Below the reference gel are the Western blots using serum pooled from groups of n=3-5 mice in the indicated groups. GPs loaded with mouse serum albumin (MSA) served as a negative vaccine control, and recombinant ovalbumin (Ova) served as a negative protein control. The blots have been cropped to show the relevant portions but have not been digitally manipulated.

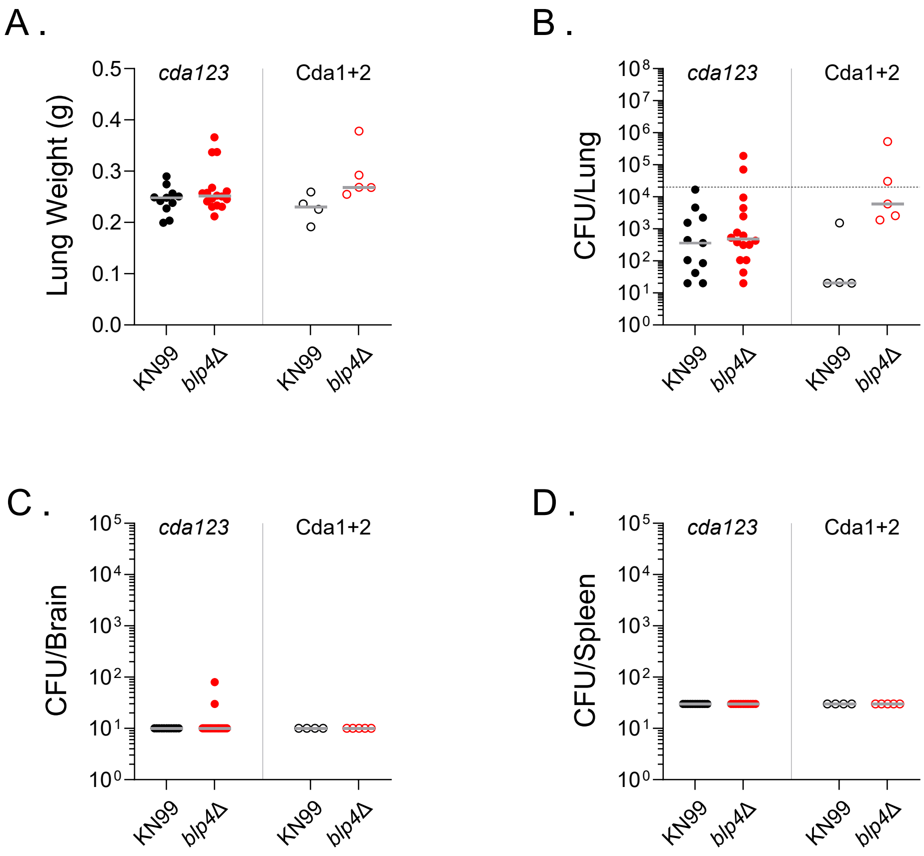


**Supplementary Figure 5: Analysis of organs from mice that survived infection with KN99 and *blp4*Δ.** BALB/c mice were vaccinated with either live *cda123* or Cda1 and Cda2 adjuvanted with CAF01 (Cda1+2) before receiving an orotracheal challenge of 2 x $10^4$ CFU of the indicated cryptococcal strain. Survival was measured until 73 days post infection. Lung weights (A) and fungal burden in the lungs (B), brains (C), and spleens (D) were determined for surviving mice. At the end of the study, surviving mice were euthanized via cardiac puncture. Dissected lungs and brains were homogenized in 4 mL, 2 mL, respectively, of PBS with 200 U/mL penicillin and 200 µg/mL streptomycin. Lungs and brains were homogenized using an OMNI TH tissue homogenizer with a tip adaptor for 7 mm X 110 mm hard tissue probe (OMNI International, Kennesaw, GA). Spleens were homogenized with a syringe piston and passed through a 70 micron filter in a total volume of 6 mL before plating. Homogenates were diluted when necessary and plated on Sabouraud Dextrose agar. CFUs were counted after two days incubation at 30°C. The dotted line on CFU/lung indicates the fungal challenge dose. The limit of detections for CFU in the lungs, brains, and spleens were 20, 10, and 30 CFU, respectively.

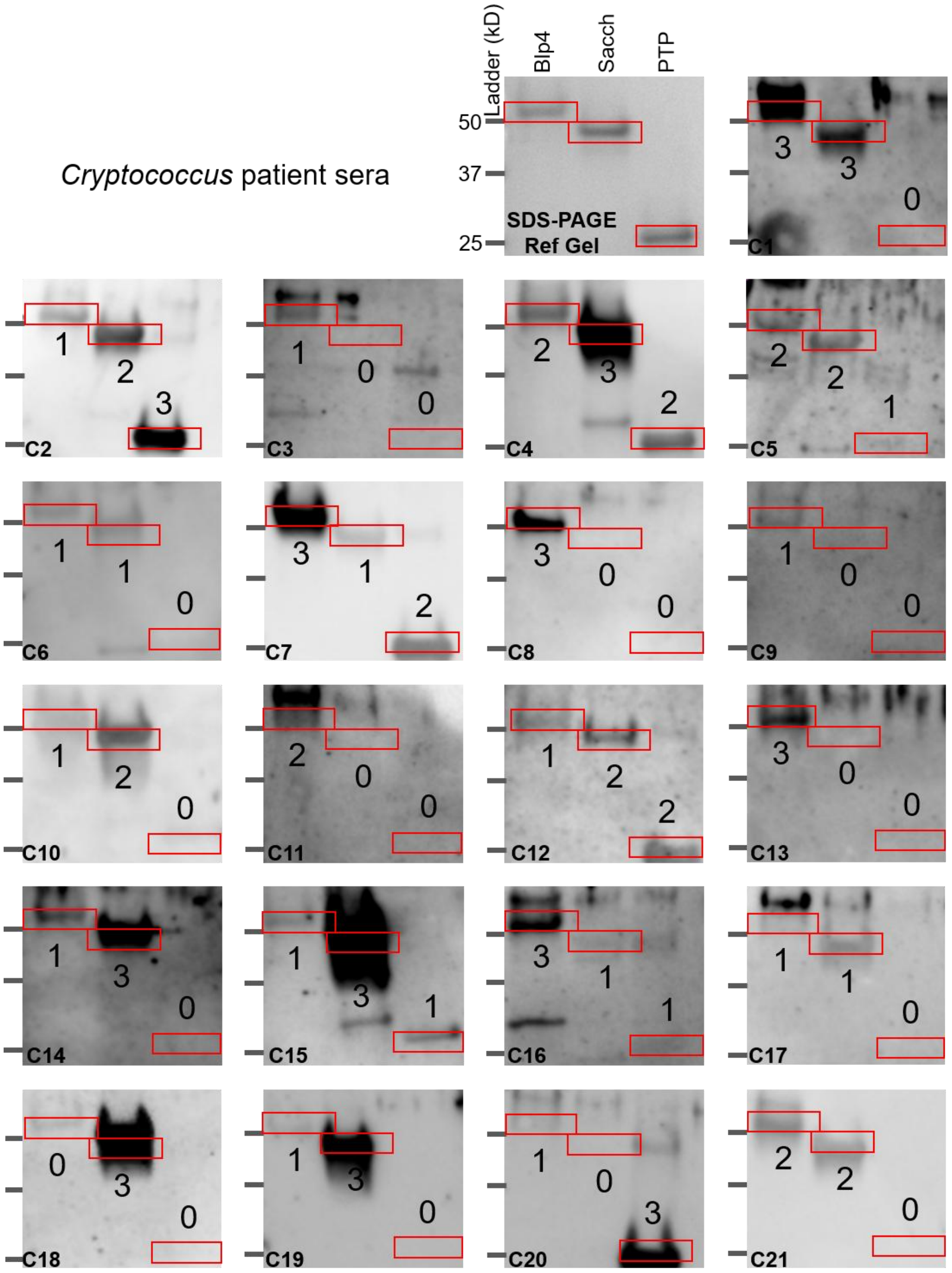


**Supplementary Figure 6: Immunoblots of *Cryptococcus* patient sera.** Individual human serum samples were probed via Western blot for antibody to Blp4, Sacch, and PTP. Protein size was visualized via SDS-PAGE (top left). Band intensity was quantified for each antigen on immunoblots from patients with a history of cryptococcosis by blind scoring by three reviewers. A score of 0 is negative, while 3 is the most positive. Red boxes indicate the correct molecular weight for the protein of interest. The median score for the 3 reviewers is denoted beneath the red box for Blp4 and Sacch, and above the red box for PTP. Sample number is denoted in the bottom left corner of each blot.

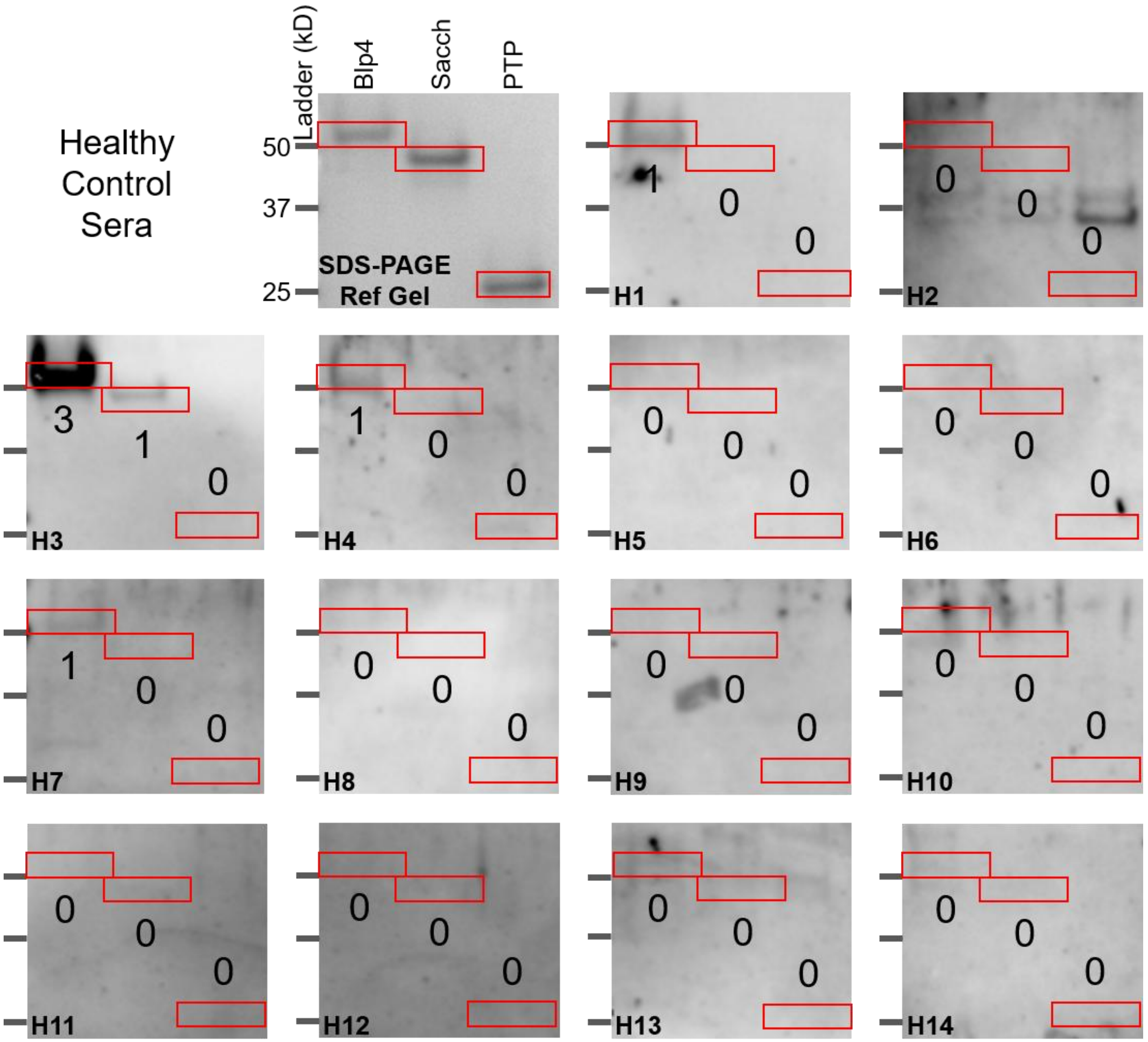


**Supplementary Figure 7: Immunoblots of healthy control sera.** Individual human serum samples were probed via Western blot for antibody to Blp4, Sacch, and PTP. Protein size was visualized via SDS-PAGE (top left). Band intensity was quantified for each antigen on immunoblots from healthy control sera by blind scoring by three reviewers. A score of 0 is negative, while 3 is the most positive. Red boxes indicate the correct molecular weight for the protein of interest. The median score for the 3 reviewers is denoted beneath the red box for Blp4 and Sacch, and above the red box for PTP. Sample number is denoted in the bottom left corner of each blot.

A.

| Sample | | Median Western Blot Scores | | | Age/ Gender | Race/ Ethnicity | Time since diagnosis | Site of cryptococcosis | *Cryptococcus* species | IgG (mg/dL) | B cell count (cells/µL) | CD4 T cell count (cells/µL) | CD8 T cell count (cells/µL) | Serum CrAg |
|---|---|---|---|---|---|---|---|---|---|---|---|---|---|---|
| | | Blp4 | Sacch | PTP | | | | | | | | | | |
| *Cryptococcus* patients | **C1*** | 3 | 3 | 0 | 62/M | W/NH | 3 years | CNS | *C. neoformans* | 434 | 88 | 594 | 519 | 1:64 |
| | **C2*** | 1 | 2 | 3 | 52/M | W/NH | 6 years | CNS | *C. neoformans* | 1,321 | 448 | 643 | 415 | Neg |
| | **C3*** | 1 | 0 | 0 | 72/M | W/NH | 1 month | CNS | *C. neoformans* | 876 | 49 | 636 | 119 | 1:4 |
| | **C4*** | 2 | 3 | 2 | 57/M | W/NH | 7 months | CNS | *C. neoformans* | 1,595 | 241 | 927 | 500 | 1:64 |
| | **C5*** | 2 | 2 | 1 | 73/F | W/NH | 38 years | pulmonary | unknown | 590 | 411 | 1,185 | 506 | Neg |
| | **C6*** | 1 | 1 | 0 | 55/F | AI/NH | 9 years | CNS | *C. neoformans* | 692 | 413 | 1,725 | 647 | Neg |
| | **C7** | 3 | 1 | 2 | 46/F | W/NH | 5 months | CNS | *C. gattii* | 485 | 228 | 986 | 405 | 1:256 |
| | **C8** | 3 | 0 | 0 | 60/M | W/NH | 6 years | CNS | *C. neoformans* | 729 | 153 | 545 | 246 | 1:32 |
| | **C9** | 1 | 0 | 0 | 70/F | BA/NH | 6 years | pulmonary | unknown | 854 | 235 | 744 | 343 | Neg |
| | **C10** | 1 | 2 | 0 | 19/F | W/NH | 2 years | pulmonary | *C. neoformans* | 1,133 | 344 | 1,331 | 785 | Neg |
| | **C11** | 2 | 0 | 0 | 6/M | W/NH | 4 years | pulmonary | *C. gattii* | 882 | 446 | 1,071 | 679 | Neg |
| | **C12** | 1 | 2 | 2 | 65/F | W/NH | 2 years | CNS | *C. neoformans* | 663 | 12 | 272 | 143 | 1:2 |
| | **C13** | 3 | 0 | 0 | 44/M | BA/NH | 9 months | CNS | *C. gattii* | 467 | 45 | 414 | 242 | 1:64 |
| | **C14** | 1 | 3 | 0 | 53/F | W/NH | 4 years | pulmonary | *C. neoformans* | 1,137 | 216 | 1,179 | 539 | Neg |
| | **C15** | 1 | 3 | 1 | 38/M | W/NH | 4 months | CNS | unknown | 560 | 107 | 851 | 284 | 1:64 |
| | **C16** | 3 | 1 | 1 | 65/F | W/NH | 3 years | CNS | unknown | 896 | 140 | 2,329 | 923 | Neg |
| | **C17** | 1 | 1 | 0 | 59/M | W/NH | 2 years | CNS | *C. neoformans* | 559 | 6,584 | 1,382 | 397 | Neg |
| | **C18** | 0 | 3 | 0 | 33/F | AAPI/NH | 1 years | CNS | *C. gattii* | 809 | 224 | 792 | 1,163 | 1:512 |
| | **C19** | 1 | 3 | 0 | 38/M | W/NH | 1 year | CNS | *C. gattii* | 334 | 137 | 1,133 | 248 | 1:128 |
| | **C20** | 1 | 0 | 3 | 43/M | AAPI/NH | 1 year | CNS | unknown | 898 | 185 | 774 | 843 | Neg |
| | **C21** | 2 | 2 | 0 | 18/F | MR/NH | 1 month | CNS | *C. gattii* | 1,072 | 104 | 453 | 184 | 1:512 |

B.

| Sample | | Median Western Blot Scores | | | Age/ Gender | Race/ Ethnicity |
|---|---|---|---|---|---|---|
| | | Blp4 | Sacch | PTP | | |
| Healthy donors | **H1*** | 1 | 0 | 0 | 51/M | BA/U |
| | **H2*** | 0 | 0 | 0 | 66/M | W/U |
| | **H3** | 3 | 1 | 0 | 51/F | W/U |
| | **H4** | 1 | 0 | 0 | U/U | U/U |
| | **H5** | 0 | 0 | 0 | U/U | U/U |
| | **H6** | 0 | 0 | 0 | U/U | U/U |
| | **H7** | 1 | 0 | 0 | 55/M | W/U |
| | **H8** | 0 | 0 | 0 | 49/F | W/U |
| | **H9** | 0 | 0 | 0 | U/U | U/U |
| | **H10** | 0 | 0 | 0 | 56/M | W/NH |
| | **H11** | 0 | 0 | 0 | U/U | U/U |
| | **H12** | 0 | 0 | 0 | U/U | U/U |
| | **H13** | 0 | 0 | 0 | U/U | U/U |
| | **H14** | 0 | 0 | 0 | U/U | U/U |

**Supplementary Table 1: Clinical and demographic information for *Cryptococcus* patient serum donors (A) and healthy control donors (B).** * indicates donor PBMCs were tested in previous *ex vivo* stimulation assays [1]. Legend for abbreviations: M: Male, F: Female, W: White, NH: Not Hispanic or Latino, AI: American Indian or Alaska Native, BA: Black/African American, AAPI: Asian American or Pacific Islander, MR: multiracial, U: Unknown, CNS: Central nervous system, CrAg: Cryptococcal antigen, Neg: negative. B cell counts were determined by CD19 expression, except for C9 where CD20 was used.
